# *Escherichia coli* ribosomal protein S1 enhances the kinetics of ribosome biogenesis and RNA decay

**DOI:** 10.1101/2021.10.20.465233

**Authors:** Mélodie Duval, Karine Prévost, Katarzyna J. Bandyra, Anne Catherine Helfer, Alexey Korepanov, Latifa Bakhti, Lauriane Kuhn, Mathias Springer, Pascale Romby, Ben F. Luisi, Eric Massé, Stefano Marzi

**Affiliations:** Architecture et Réactivité de l’ARN, Université de Strasbourg, IBMC-CNRS, F-67084 Strasbourg, France; Université de Sherbrooke, CRCHUS, Faculty of Medicine and Health Sciences, Department of Biochemistry and Functional Genomics, RNA Group, 3201 Jean Mignault Street, Sherbrooke, Quebec, Canada, J1E 4K8; Institut de Biologie Physico-Chimique CNRS, 75005; Department of Biochemistry, University of Cambridge, Tennis Court Road, Cambridge CB2 1GA, UK; Plateforme protéomique Strasbourg-Esplanade, IBMC-CNRS, Strasbourg, France

**Keywords:** ribosomal protein S1, sRNA-dependent regulation, mRNA turnover, rRNA maturation, RNA chaperone

## Abstract

*Escherichia coli* ribosomal protein S1 is essential for translation initiation of mRNAs and for cellular viability. Two oligonucleotide binding (OB)-fold domains located in the C-terminus of S1 are dispensable for growth, but their deletion causes a cold-shock phenotype, loss of motility and deregulation of RNA mediated stress responses. Surprisingly, the expression of the small regulatory RNA RyhB and one of its repressed target mRNA, *sodB*, are enhanced in the mutant strain lacking the two OB domains. Using *in vivo* and *in vitro* approaches, we show that RyhB retains its capacity to repress translation of target mRNAs in the mutant strain but becomes deficient in triggering rapid turnover of those transcripts. In addition, the mutant is defective in of the final step of the RNase E-dependent maturation of the 16S rRNA. This work unveils an unexpected function of S1 in facilitating ribosome biogenesis and RyhB-dependent mRNA decay mediated by the RNA degradosome. Through its RNA chaperone activity, S1 participates to the coupling between ribosome biogenesis, translation, and RNA decay.

## Introduction

Translation initiation is the rate limiting step of protein synthesis and is regulated in various ways throughout all domains of life (1–7). In bacteria, many messenger RNAs (mRNAs) carry regulatory elements that directly sense the environmental cues or that are specifically recognized by a variety of trans-acting ligands (sRNAs, RNA-binding proteins) to regulate translation initiation. Some of these regulatory elements are characterized by structures that potentially interfere with ribosome recognition. *E. coli* ribosomal protein S1 (r-protein S1) is one of the key proteins involved in translation initiation, primarily through its action to help recruit and correctly position mRNAs carrying structured 5’UTRs or/and with suboptimal Shine and Dalgarno (SD) sequences (7–11). Several studies have shown that S1 contributes an RNA melting activity (12–16) that may facilitate the early steps of translation initiation (7). Besides its critical role in translation, the r-protein S1 has been implicated in other cellular processes, such as transcription recycling (17), rescuing of stalled ribosomes by tmRNA (18–21), and repressing its own expression (22). In cooperation with r-protein S2, it inhibits the translation of *rpsB* mRNA (23). Overproduction of S1 stabilizes *pnp* mRNA, encoding the exoribonuclease polynucleotide phosphorylase(24), as well as protection of specific mRNAs against RNase E attack (25). Finally, r-protein S1 is also part of various multi-protein complexes, one of which assists the degradation of mRNAs by RegB endoribonuclease (26) while another is required for replication of the Qβ phage (27–30).

In *E. coli*, S1 protein is composed of 6 oligonucleotide binding (OB)-fold domains, which are conserved among Gram-negative bacteria and few Gram-positive bacteria (31). The N-terminal domains 1 and 2 are responsible for ribosome binding (7, 32–35), and the minimal r-protein S1 that is required for translation initiation of many mRNAs is composed of domains 1 to 4 (7, 22, 28). These four domains are also essential for cell viability (7). However, the role of domains 5 and 6 of S1 remains less clear. Domain 5 is almost identical to domain 4 and reinforces the RNA binding capacity (7, 28, 31, 32), and domain 6 may participate in the recycling of RNA polymerase (17). A phylogenetic analysis revealed that this C-terminal domain is conserved in *Enterobacteriaceae* (31). The deletion of the last two C-terminal domains of *E. coli* S1 led to a viable mutant strain, albeit with a slower growth and a cold-sensitive phenotype (7, 36).

In this work, we have addressed the functions of the last two domains of *E. coli* r-protein S1. Comparative proteomic and RNA-seq analysis performed on the wild-type strain and the mutant strains depleted of domain 6 (*rpsA*Δ6), or of both domains 5 and 6 (*rpsA*Δ56) revealed an increased expression of a large set of genes responding to various stresses, and a reduced expression of most motility genes. Moreover, the expression of many sRNAs was more abundant in the mutant strains such as RyhB, which is one of the best characterized sRNA in *E. coli*. Strikingly, the expression of many of the RyhB-repressed targets was also enhanced in the mutant strains despite the higher levels of that sRNA. Remarkably, although the effect of RyhB on mRNA binding and translation was not altered, there was strong impairment in degradation of the repressed mRNAs by the RNA degradosome. Furthermore, our results also showed that the 16S rRNA maturation was slower in the absence of both domains 5 and 6 of S1. We describe a functional link between the C-terminal domains of S1 and the RNA degradosome and that kinetic of mRNA degradation and rRNA maturation assisted by r-protein S1 is an important feature for bacterial fitness.

## Results

### Deletion of the last two domains of S1 deregulates regulatory RNAs and genes for stress responses and motility

Various mutant strains were previously constructed where the OB-fold domains were successively depleted from the C-terminus (**Figure S1A**) (7). The mutant strains were sequenced to verify that no additional mutation, acting as suppressors, had occurred. The only viable mutants were those deleted for domain 6 alone (*rpsA*Δ6), and for domains 5 and 6 together (*rpsA*Δ56) even though they harbored an increased doubling time and a longer lag phase than the WT strain (**Figure S1B**). Phenotypic assays were performed to monitor the motility of the WT and mutant strains (**Figure S1C**). Using semisolid agar plates to measure the characteristic chemotactic rings of the bacterial colonies produced by bacterial swarming, the *rpsA*Δ6 and *rpsA*Δ56 mutant strains showed less colony spreading, reflecting defect in motility. We then measured single cell swimming ability by tracking movements of several bacteria in liquid medium under the microscope for the WT and *rpsA*Δ56 strains (**Figure S1D**). For the WT strain, a classical behavior was observed with tracks corresponding to both swimming and tumbling, while in striking contrast the *rpsA*Δ56 mutant strain reproducibly remained motionless.

Because both mutant strains have similar phenotypic behaviors, we analyzed the effect of the deletion of the two last C-terminal domains of S1 on gene expression using differential transcriptomics and proteomics. Label-free mass spectrometry performed on the *rpsA*Δ56 mutant and the isogenic WT strains identified 342 proteins, which were significantly altered in relative abundance in the mutant strain (threshold 2-fold, p-values ≤0.05, **Figure 1A** and **Table S1**), representing 23% of the total detected proteins (1509). Using RNA-Seq, 835 RNAs were identified with altered expression in *rpsA*Δ56 mutant strain (threshold 2-fold, p-values ≤0.05, **Figure 1B** and **Table S2**), representing 19% of the detected transcripts (4442). The two approaches were particularly well correlated for genes encoding proteins involved in motility (FliC, FliG, FlhC) and chemotaxis (CheZ, CheR, CheA) (**Table 1**). Indeed, the decreased yields of these mRNAs were accompanied by a strong drop of the levels of the corresponding proteins in the *rpsA*Δ56 mutant strain.

**Figure 1:**
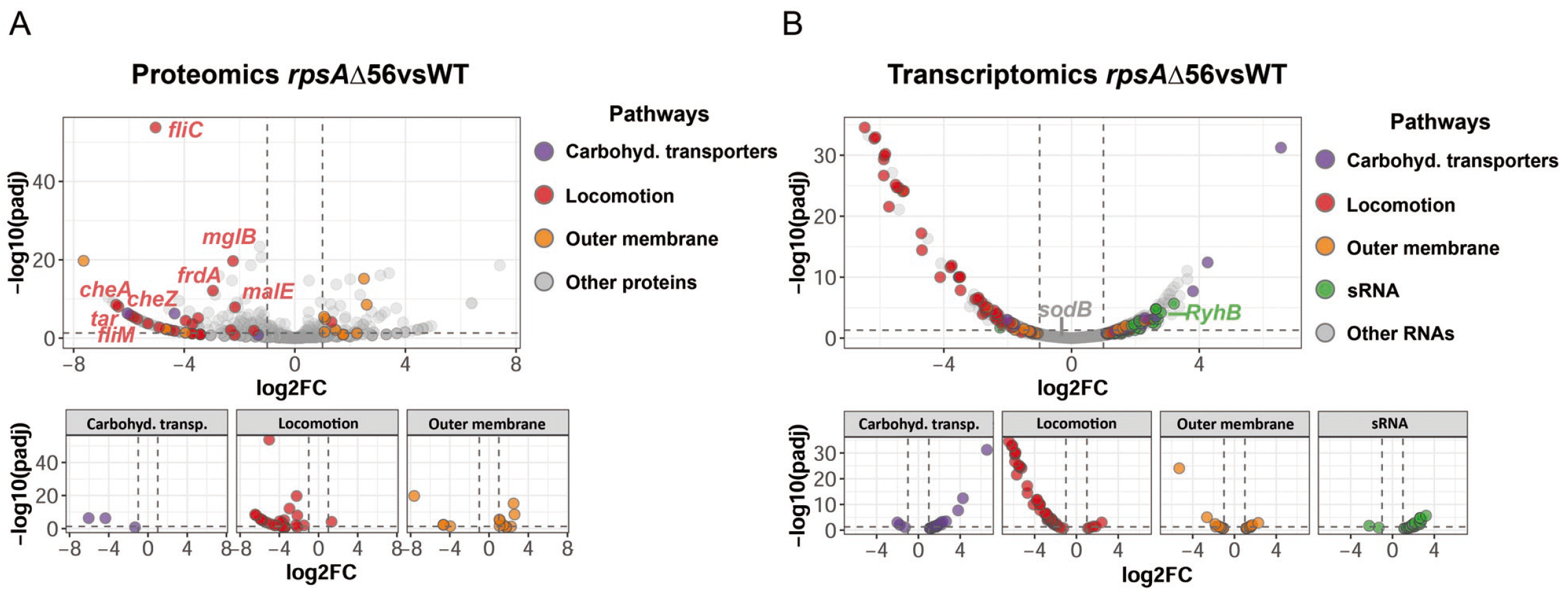
Comparison of gene expression in wild-type (WT) and mutant strains by proteomic and RNA-seq analysis. (A) Comparative proteomic analysis of the proteins expressed in the WT and in the *rpsA*Δ56 strain. The threshold was set at an induction fold of 2 (P-value <0.05). Several proteins are colored according to the metabolic and functional pathways to which they belong. (B) Comparative RNA-seq analysis of the RNA expressed in the WT and in the *rpsA*Δ56 strain. The threshold was set at an induction fold of 2 (P-value <0.05). The raw data are provided in Table S1 (proteomic analysis WT vs *rpsA*Δ56 strains) and Table S2 (RNA-seq analysis WT vs *rpsA*Δ56 strains).

**Table 1:**
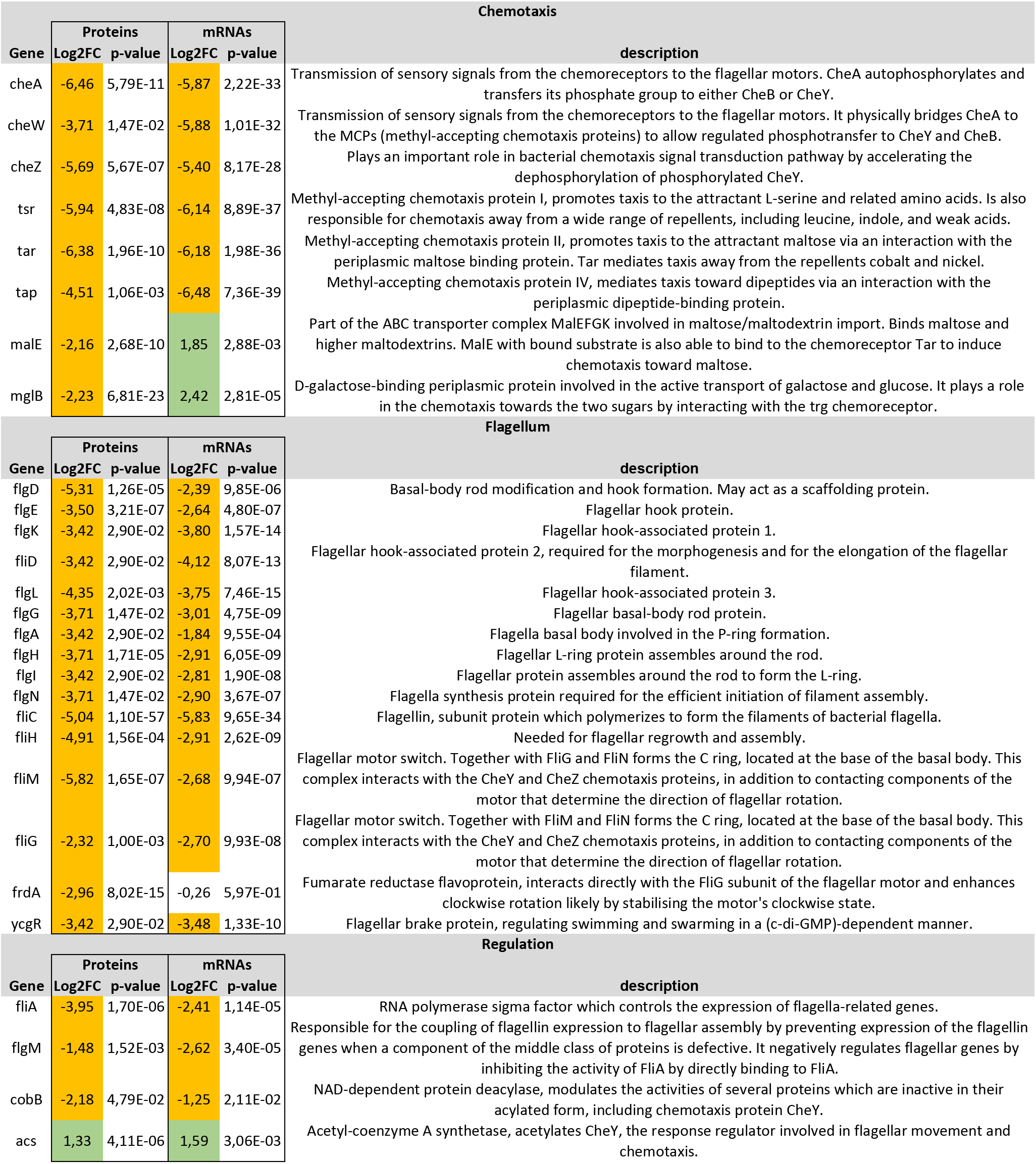
Comparative proteomics and transcriptomics analysis performed on the WT and *rpsA*Δ56 mutant strains. The data are given for genes encoding proteins that are involved in chemotaxis and motility.

The steady state levels of a significant number of mRNAs involved in various stress responses were enhanced in the *rpsAΔ56* mutant strain. These mRNAs encoded proteins that are involved in heat shock, osmotic stress, iron metabolism, oxidative stress, and SOS responses (**Figure 1B** and **Table S2**). In addition, the level of two sRNAs was lower while the expression of 30 sRNAs was slightly enhanced in the mutant strain (**Figure 1B** and **Table S2**). Among these sRNAs, RyhB was the second most upregulated sRNA (**Table S2**). Since RyhB-dependent repression is often associated with rapid depletion of the mRNA targets, we analyzed more precisely the expression of the known RyhB-dependent targets (**Table S3**). However, the comparison between transcriptomic and proteomic analysis in WT and *rpsAΔ*56 mutant strains revealed complex responses (**Table S3**). The yields of few mRNAs (*flgA*, *cydA*, *cra*) were slightly decreased in the mutant strain accompanied with a decreased of the protein levels, in agreement with higher yields of RyhB in the mutant strain. However, several mRNA levels remained unchanged (i.e., *frdA*, *iscA*, *fumA*, *bfr, nuoF*) or were slightly enhanced (*sdhB*) while the protein yields were significantly reduced in the mutant strain (**Table S3**). The fact that these RyhB-dependent mRNA targets are not degraded in the mutant strain is not attributable to a decreased level of RNase E or PNPase, which are similar to those of the WT strain (**Table S1**). These data suggested that the full length S1 protein might be required for rapid depletion of mRNAs targeted for repression by some sRNAs.

We next explored the possible action of S1 in RyhB-dependent regulation under conditions where RyhB exertes its regulatory functions, i.e., depletion of iron.

### RyhB-dependent *sodB* repression is altered in the mutant strain

Because S1 is dispensable for *sodB* translation initiation (7), we first analyzed the role of S1 in RyhB-dependent regulation of *sodB*. The effect of RyhB expression on *sodB* mRNA levels was monitored in the WT and mutant *rpsA*Δ56 strains. After purification of total RNAs at several time points, Northern blot assays were performed with probes complementary to either *sodB* or RyhB (**Figure 2A**). As a loading control, we probed for 5S rRNA (**Figure S2A**). As described previously, RyhB expression was induced by the addition of the iron chelator 2,2ʹ-dipyridyl (Dip.) in the medium (37) (**Figure 2A**). After 5 min, the medium was supplemented with sufficient iron sulfate (FeSO_4_) to inhibit RyhB synthesis. As expected, in the WT strain, *sodB* levels dropped immediately upon RyhB induction and were restored upon the addition of iron. In contrast,, high levels of *sodB* were constantly observed in the *rpsA*Δ56 mutant strain whatever the induction or the repression of RyhB synthesis. Upon addition of iron, RyhB remained detectable for a longer time in the *rpsA*Δ56 mutant strain than in the WT strain, in agreement with the transcriptomic analysis (**Table S2**). These data showed that the lack of domains 5 and 6 in S1 influenced the levels of both RyhB sRNA and *sodB* mRNA.

**Figure 2:**
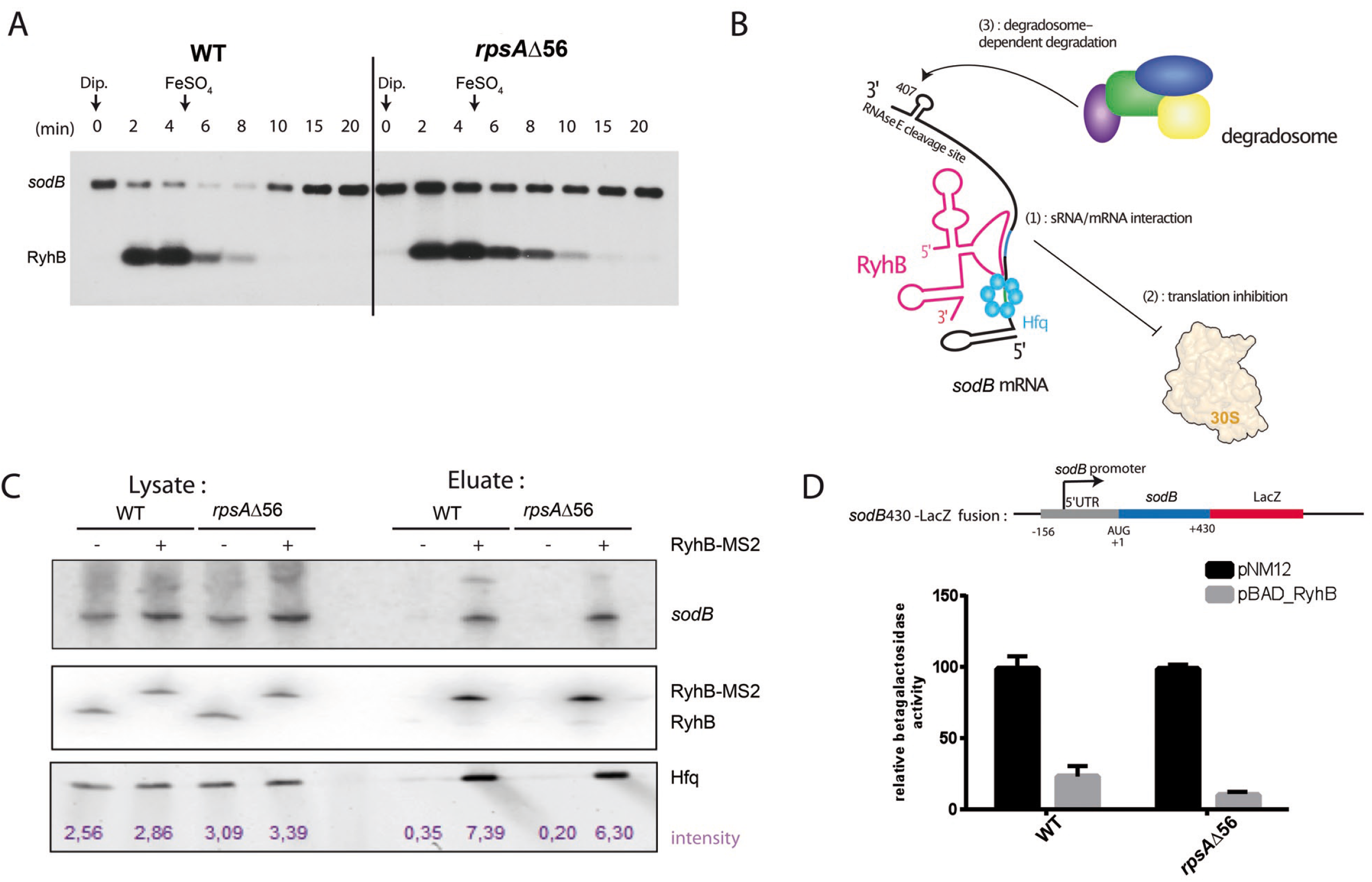
The deletion of S1 C-terminal domains perturbs RyhB-mediated *sodB* degradation. (A) Northern blot analysis was performed using a labeled probe against either *sodB* or RyhB. Total RNA extracts were prepared from WT and *rpsA*Δ56 cultures in LB at 37°C. At OD_600_=0.4, 250 μM 2,2’-dipyridyl was added to the culture to induce RyhB. After 5 min, 100 μM of FeSO_4_ was added to specifically inhibit RyhB synthesis. The same samples were run on another gel for 5S rRNA (5S) detection, as a loading control. (B) RyhB-mediated *sodB* regulation has been well described, and occurs in three steps: (1) sRNA-mRNA form a complex together with the protein Hfq, (2) RyhB inhibits translation initiation, and induces (3) rapid degradation of the sRNA/mRNA duplex by the RNA degradosome. (C) Northern Blot analysis performed on RNA crude extracts prepared from WT and mutant *rpsA*Δ56 strains expressing RyhB fused with the MS2 tag (38) and purified on affinity chromatography. The presence of *sodB* and RyhB was monitored with appropriate labeled probes, and anti-FLAG antibodies were used for Hfq^3xFLAG^ Western Blot analysis. The synthesis of MS2-RyhB (+) and RyhB (control) expressed from a pBAD plasmid, was induced by the addition of 0.1 % arabinose in the indicated lanes. (D) Analysis of ß-galactosidase synthesis from the *sodB_430_-lacZ* translational fusion integrated into the chromosome of the MG1655 (WT) and mutant (*rpsA*Δ56) strains in presence or absence of RyhB. The *lacZ* gene was fused to *sodB* containing 430 nucleotides of its coding region from the AUG including the major RNase E cleavage site. Strains carried either an empty vector (pNM12; black) or a pBAD-*ryhB* (grey). R*yhB* expression was induced by addition of 0.1% arabinose. Signals from each strain were normalized according to the corresponding empty vector. Data are representative of three independent experiments.

It was previously demonstrated that RyhB-dependent *sodB* degradation occurred in three main steps (**Figure 2B**): (1) formation of sRNA-mRNA binding together with the protein Hfq, (2) translation inhibition followed by (3) the subsequent rapid degradation of the sRNA/mRNA duplex by the RNA degradosome (38). Because the rapid depletion of the mRNA is a consequence of the repression of translation, we analyzed if S1 might be required for RyhB binding and translation repression. First, we used MS2-sRNA affinity purification coupled to Northern Blot to probe the interaction between MS2-RyhB and *sodB* mRNA in the WT and mutant *rpsA*Δ56 strains (39) (**Figure 2C**). MS2-RyhB construct was expressed from a plasmid (pBAD-MS2-RyhB, **Table S4**) *via* arabinose induction in WT and mutant *rpsA*Δ56, and the crude extract was loaded on an affinity matrix containing the maltose binding protein fused to MS2 protein. As negative control, untagged RyhB was expressed in the same conditions from the pBAD-RyhB plasmid (**Table S4**). Northern Blot experiments were carried out to visualize RyhB and *sodB* in the lysate and eluate fractions. The data revealed that *sodB* was specifically retained in fractions containing MS2-RyhB and that the same amount of *sodB* was detected in the WT and *rpsA*Δ56 strains. Western Blot analysis showed that Hfq was also bound to the duplex at a similar level in WT and *rpsA*Δ56 strains. Overall, these data showed that the formation of base-pairing interactions between *sodB* and RyhB is not perturbed by the deletion of the C-terminal domains of S1 *in vivo*.

We then assessed the ability of RyhB to repress *sodB* translation in the WT and mutant strains. A *sodB-lacZ* reporter fusion under the control of the endogenous *sodB* promoter was integrated into the chromosome of both strains and the activity of β-galactosidase was measured (**Figure 2D**). The synthesis of RyhB expressed from a plasmid under the control of an inducible promoter caused a strong decrease of the β-galactosidase synthesis in both the WT and in the *rpsA*Δ56 strains. We then analyzed the ability of RyhB to repress the translation of *sodB* using *in vitro* translation assays supplemented with the ribosomes purified from the WT and *rpsA*Δ56 strains. Ribosomes from the mutant and parental strains were able to translate *sodB* with the same efficiency (**Figure S2B**). The incorporation of the S^35^Met showed that the addition of increasing concentrations of RyhB reduced considerably the synthesis of SodB protein with the two sets of ribosomes (**Figure S2C**). Finally, toeprinting assays were used to monitor the effect of RyhB binding on the formation of the ternary initiation complex involving the 30S ribosomal subunits containing WT S1 or the truncated protein S1Δ56, the initiator tRNA^Met^, and *sodB* mRNA. We have verified the quality of the 30S purification using mass spectrometry analysis (results not shown). Formation of the initiation complex blocks the elongation of a cDNA primer by reverse transcriptase and induces a signal at position +16 (the A of the initiation codon being the +1, **Figure S2D**). Binding of RyhB to *sodB* mRNA strongly decreased the formation of the active initiation complex whatever the nature of S1 present on the 30S subunits.

Taken together, these data showed that RyhB can repress *sodB* translation with the same efficiency in the WT and *rpsA*Δ56 strains.

### S1 is required for the rapid RyhB-dependent depletion of *sodB* mRNA

We then assessed whether S1 is involved in RyhB-dependent degradation of *sodB* mediated by the RNA degradosome. RyhB expression was induced by the addition of 2,2’-dipyridyl during a long period of 25 min, during which levels of *sodB* and RyhB were evaluated. Northern Blot analysis showed that *sodB* remained detectable for a longer period in the *rpsA*Δ56 mutant strain than in the WT, indicating that the degradation of *sodB* occurred with a slower kinetic in the mutant strain due to the absence of domains 5 and 6 of S1 (**Figure 3A**). We then measured the half-life of *sodB* mRNA upon RyhB expression in both strains by adding rifampicin. The results showed that *sodB* is degraded over ten-fold faster in the WT strain (less than 1 min) than in the *rpsA*Δ56 mutant strain (11 min) (**Figure 3B**).

**Figure 3:**
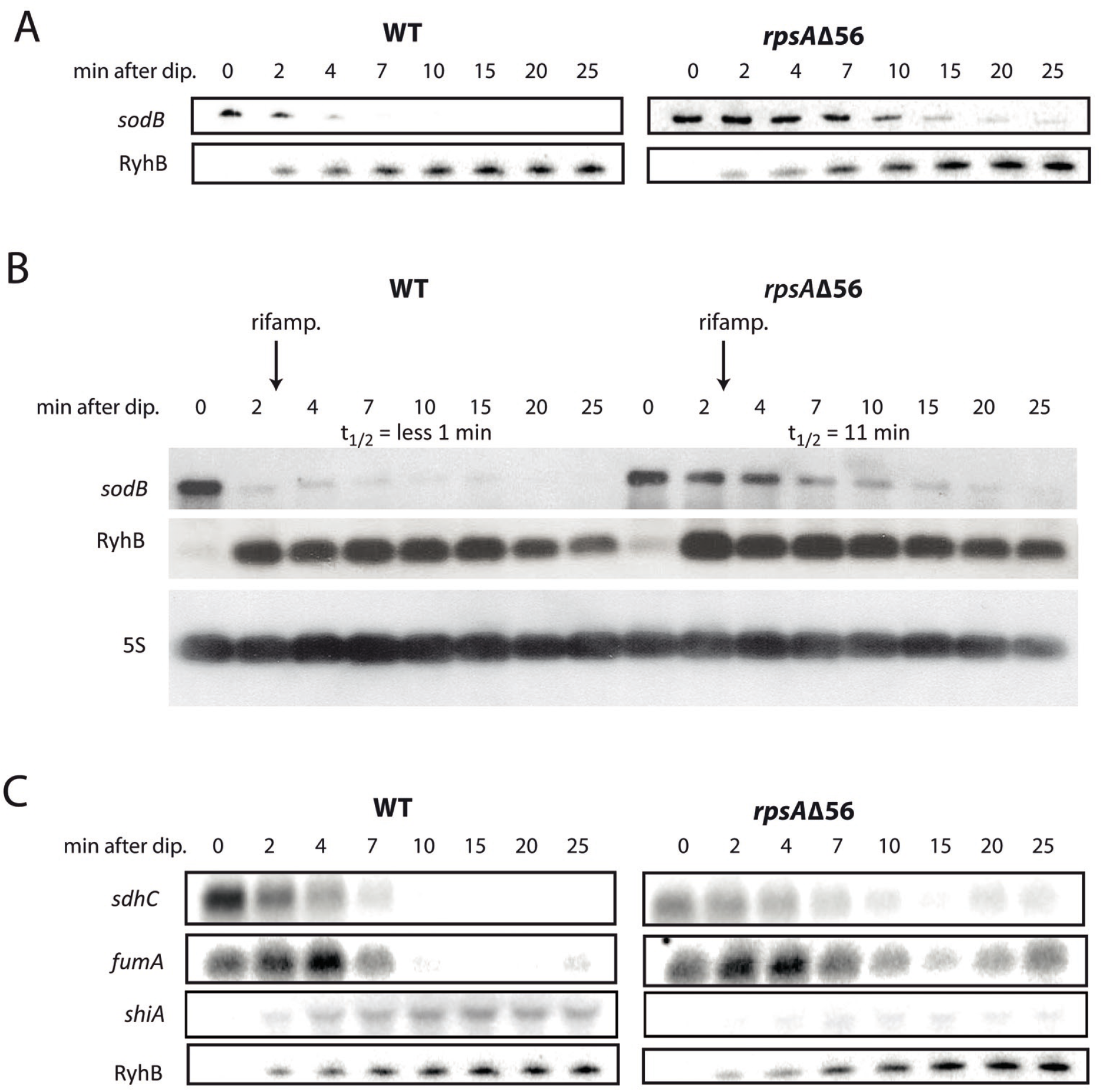
The RNA degradosome requires the C-terminal domains of S1 to induce fast RyhB-mediated degradation of *sodB*. (A) Same legend as in Figure 2A, except that FeSO_4_ was not added to the culture to visualize longer degradation pattern. The results show that in the absence of the last two domains of S1, *sodB* mRNA is degraded in a much slower manner in response to RyhB induction than in the WT strain. (B) Measurements of the half-life of *sodB* mRNA in the WT and *rpsA*Δ56 mutant strains. The experiment was done by adding rifampicin at 2 min (2,2`-dipyridyl induction being the t=0), to block the transcription and to visualize the degradation of *sodB*. t1/2 represents the half-life which was derived after quantification of the autoradiographies. RyhB and 5S were detected using the same RNA samples, which were run on different gels in parallel. (C) Analysis of the mRNAs, which belong to the RyhB regulon. Same legend as in Figure 3A.

RyhB induces the rapid degradation of more than 17 mRNAs encoding non-essential Fe containing proteins such as *sodB*, *fumA*, *sdhCDAB*, *iscA*, and *erpA* (39–44). In addition, RyhB activates the translation of *shiA* mRNA by disrupting its inhibitory secondary structure (45). In order to assess whether the effect of the absence of the last two domains of S1 on *sodB* can be generalized to other RyhB targets, Northern blot analysis was performed under conditions where RyhB expression was induced. As expected, the yields of *sdhC* and *fumA* mRNAs rapidly decreased upon the induction of RyhB (below 7 min) in the WT strain. Concomitantly, the levels of *shiA* mRNA were enhanced after 4 min of 2,2’-dipyridyl treatment (**Figure 3C**). The same experiment performed with the mutant strain showed that the rapid depletion of *sdhC* and *fumA* was altered in a manner analogous to *sodB*. More surprisingly, *shiA* mRNA was poorly detectable even after a prolonged expression of RyhB (**Figure 3C**). These data strongly suggested that the deletion of the last two domains of S1 alters the kinetics of the turnover of the RyhB-dependent mRNA targets.

We then analyzed whether the effect of S1 on mRNA degradation can be recapitulated *in vitro*. We first purified the ribosomes from the WT and *rpsA*Δ56 mutant strains and confirmed with mass spectrometry that all r-proteins were present in both ribosome preparations (result not shown). Complex between uniformly radiolabeled *sodB* mRNA and RyhB was pre-formed. The purified RNA degradosome was added either to the WT ribosomes (70S WT) containing the full length S1, to the WT ribosomes from which S1 was removed before the experiment (70S WT -S1), or to the *rpsA*Δ56 ribosomes (70S Δ56). Quantification of the full-length *sodB* mRNA at different time points showed that its degradation was reproducibly slower in the presence of ribosomes purified from *rpsA*Δ56 mutant strain than with the WT ribosomes (**Figures 4A and S3A**). Surprisingly, WT ribosomes depleted of S1 behaves as the WT ribosome containing S1, and the addition of purified S1 and S1Δ56 proteins in the absence of ribosomes had no major effect on the activity of the RNA degradosome on *sodB* mRNA *in vitro* (**Figure S3B**). Hence, these data suggest that the mutant 70S Δ56 ribosomes might be responsible for the *in vitro* slower degradation of *sodB*.

**Figure 4:**
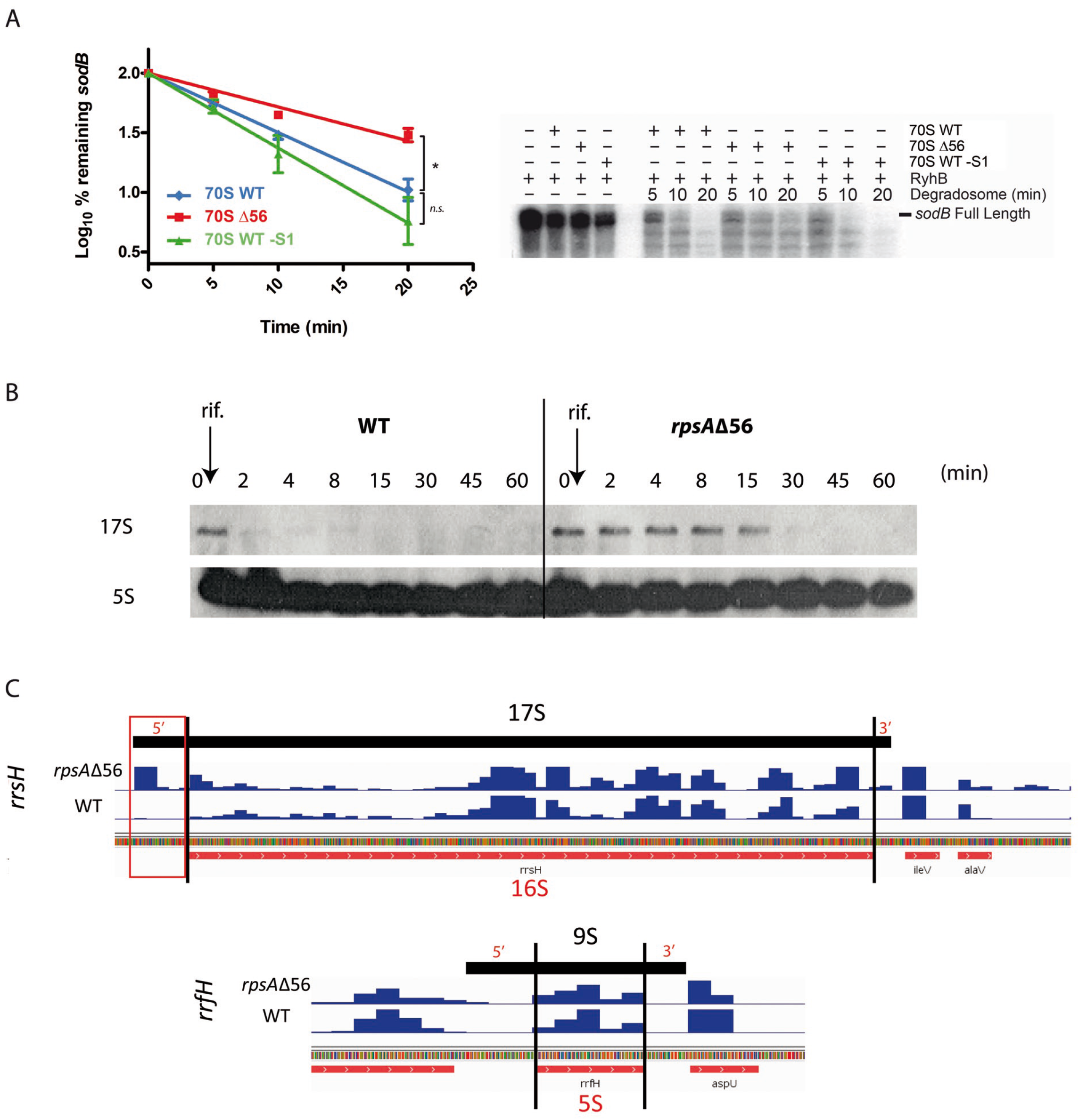
The last two domains of S1 are required for 16S rRNA maturation but not for 5S maturation. (A) *In vitro* reconstitution of *sodB* degradation using the purified RNA degradosome. Uniformly radiolabeled *sodB* mRNA was incubated in presence of RyhB, the purified RNA degradosome, and either the WT ribosome containing full length S1, the WT ribosome from which S1 was removed before the experiment, or the *rpsA*Δ56 ribosomes. The degradation of *sodB* mRNA was followed over time (5, 10, 20 min) and the RNA fragments were fractionated on an 8% polyacrylamide-7 M urea gel electrophoresis. The signals for the remaining full length *sodB* mRNA were quantified with ImageQuant TL software (GE Healthcare Life Sciences) on three independent experiments to calculate the error bars. (B) Measurements of the half-life of the 17S rRNA precursor using Northern Blot analysis. Total RNA was extracted from the WT and mutant *rpsA*Δ56 strains at various time points after addition of rifampicin. A specific probe revealed the 5’ region of the 17S. (C) IGV visualization of the rRNA reads obtained by ribosome profiling performed in WT and *rpsA*Δ56 strains. Accumulation of reads corresponding to 5’ leader of 17S precursor is observed in the mutant strain. We showed the data on *rrsH* gene as representative of the 7 rRNA operons. In comparison, reads aligned to the 5S precursor region (9S) are shown (*rrfH* gene).

### Deletion of the C-terminal domains of r-protein S1 impacts 16S rRNA maturation

The activity of the components of the RNA degradosome, and especially RNase E and PNPase, are not restricted to mRNA degradation. They are important players in other pathways such as rRNA maturation (46). Given that the mutant strain harbored a cold-sensitive phenotype, and that 70S purified from this strain affected the *in vitro* kinetics of *sodB* degradation by the degradosme, we investigated whether the absence of the last two domains of S1 might perturb the RNA degradosome activity in rRNA biogenesis.

A significant amount of 17S rRNA precursor was found in *rpsA*Δ56 mutant strain using total RNA prepared from rifampicin treated cultures and visualized on ethidium bromide stained agarose gel (**Figure S3C**). Similar data were observed using Northern Blot analyzed with a specific oligonucleotide probe complementary to the 5’ end of 16S rRNA region, which is normally cleaved by RNase E (**Figure 4B**). The results showed that the precursor is observed during a longer time period in the *rpsA*Δ56 strain (≥ 15 min) than in the WT strain (< 2 min), suggesting that the 17S precursor is processed more slowly in the absence of the last two C-terminal domains of S1. We then performed sequencing on RNA samples prepared from polysome preparation purified from the WT and *rpsA*Δ56 mutant strains. In agreement with the previous data, accumulation of reads was observed upstream of 16S gene only in the mutant strain. Interestingly, analysis of the 5S rRNA locus, which is matured from the 9S transcript by RNase E (47), did not show any accumulation of reads in the mutant strain (**Figure 4C**).

These data suggested that fast kinetics of the 5’ end of 16S rRNA processing mediated by the RNA degradosome requires the full-length r-protein S1.

## Discussion

The present study highlights unexpected features of r-protein S1 in sRNA regulation and rRNA maturation. First, we have demonstrated that the deletion of the last two C-terminal domains of S1 impairs cell motility and causes stress responses. This is accompanied by the deregulation of several genes including many sRNAs. Second, we showed that these domains on S1 are required for the rapid depletion of RyhB-dependent repressed mRNAs. Third, we demonstrated that the full-length S1 is required for normal 16S rRNA maturation.

The major functions of S1 are linked to the ribosome where the protein occupies a strategic position at the junction of the platform and the body of the 30S subunit at the solvent side close to S2 r-protein (32). Its six OB fold RNA binding domains confer to S1 the ability to recognize many mRNA substrates and to capture them at unpaired AU-rich sequences, primarily located upstream the SD sequence. S1 is essential for the recruitment and accommodation of mRNAs characterized by structured elements within the ribosome binding sites, and which contain a weak SD sequence (7, 48, 49). Thanks to its RNA chaperone activity, the protein remodels structured RNA elements in a step-wise manner to shift the structured mRNAs from a stand-by position to its accommodation into the decoding center (7, 49, 50). Besides this essential role in translation initiation, *E. coli* r-protein S1 can also act without the ribosome, free or in complex with other proteins, (1) to regulate the translation of specific mRNAs, (2) to protect mRNAs against the degradation by RNase E and the RNA degradosome, and (3) to provide additional RNA binding capacity to other protein partners (reviewed in (34)). Finally, many translational repressors (protein or sRNA) target directly the S1 functioning on the ribosome to prevent the formation of the initiation complex (7, 48, 49).

*E. coli* cold-sensitive phenotype, as the one observed for *rpsA*Δ56 mutant strain, has been previously associated with ribosome maturation defects (51). This phenotype was for instance observed for several deletion mutants of ribosome biogenesis factors (52–54) and more recently for the deletion mutant of the RNA chaperone protein Hfq (55). We showed here that the deletion of the last two C-terminal domains of the r-protein S1 affected the kinetics of the maturation of the 5’ end of 16S rRNA, which is performed by RNase E. Ribosomal protein S1 is the last protein incorporated into the ribosome, concomitant with the maturation of 17S into 16S rRNA, which occurs as the latest event of rRNA biogenesis. Due to its RNA chaperone activity, the protein might be required to modify the 17S rRNA structure that could otherwise slow down the RNase E accessibility and activity (**Figure 5A, upper panel**). Due to its localization close to the exit site of the 30S subunit, it is tempting to propose that the six OB-fold domains are necessary to reach the 5’ end of the 16S rRNA. In other words, S1Δ56 would be too short to attain the maturation site. Ribosome biogenesis is a complex process that occurs co-translationally and involves numerous factors in a well-defined orchestrated scenario where S1 would contribute to the efficient and complete biogenesis of the ribosome. Interestingly, such a role of S1 in the regulation of ribosomal biosynthesis has been proposed in *Shewanella oneidensis*, a γ-proteobacterium where the 6 OB-fold domains of S1 are similar to *E. coli* (56).

**Figure 5:**
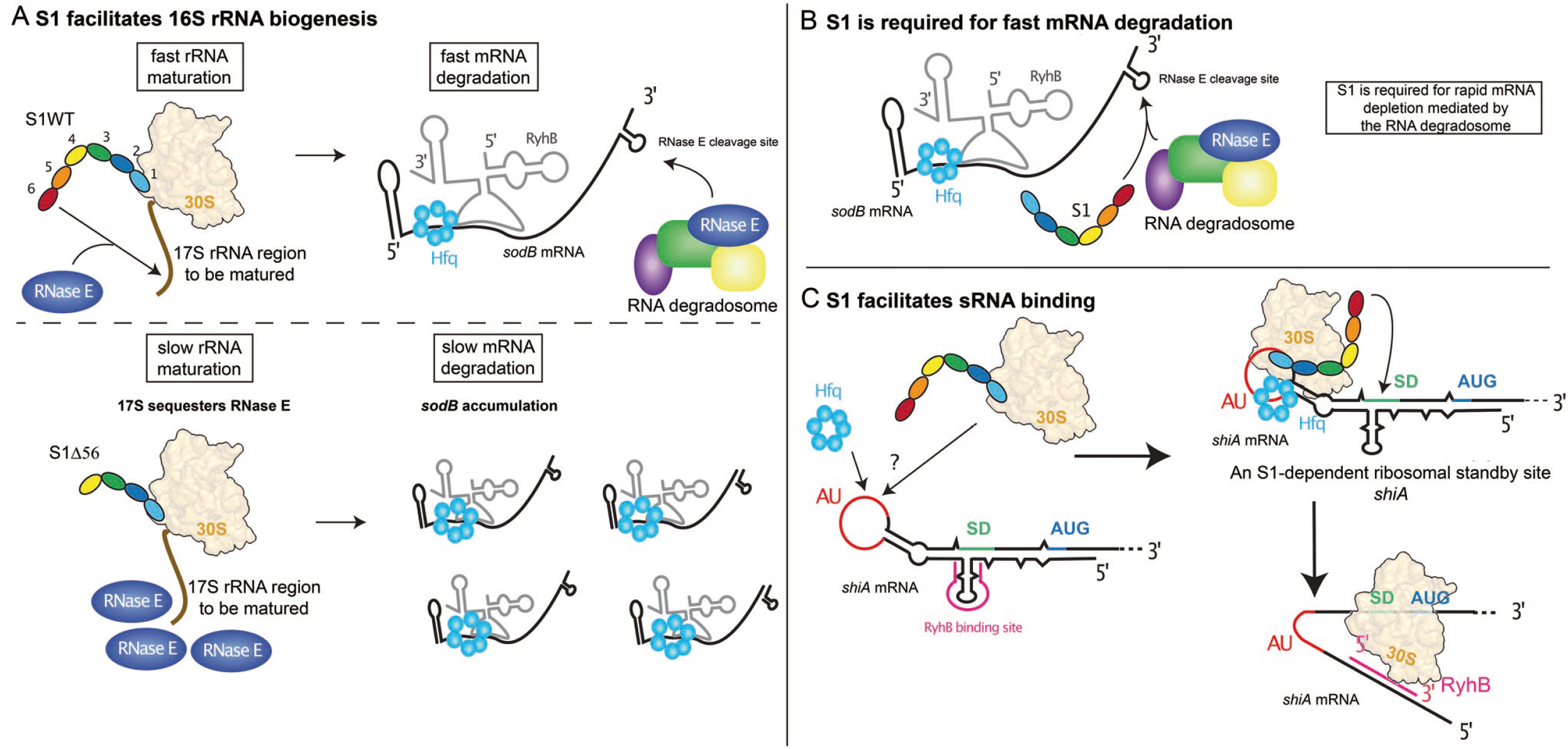
Possible models for S1 action. (A) Model for an indirect effect of S1 on *sodB* degradation. (left) S1 enhances the kinetics of 17S rRNA maturation. In WT strain, RyhB-dependent degradation of *sodB* is efficient. In the mutant *rpsA*Δ56 strain, the 17S is not efficiently matured, and it is proposed that RNase E might remain associated for a longer time with the 17S to cleave it. As the results, the kinetics of the degradation of *sodB* mRNA is altered. (B) Alternative model for direct effect of S1 on *sodB* degradation. As Hfq, S1 might be a partner to recruit the RNA degradosome at the proper site of the target mRNA. (C) A possible role of S1 in the activation of translation of *shiA* mRNA. RyhB together with Hfq favors the recruitment of the initiation ribosomal complex, which in turn stabilizes the mRNA. An unpaired AU rich sequence upstream the SD sequence might be the binding site for S1. This binding would facilitate the recruitment of the ribosome.

The deletion of the S1 C-terminal domains surprisingly causes a significant increase in the steady-state levels of many sRNAs, and especially RyhB. We showed that in *rpsA*Δ56, many mRNAs that belong to RyhB regulon remained more stable over time when the expression of RyhB is induced under iron depletion. Our data implies a functional link between S1 and the RNA degradosome as the protein enhanced the kinetics of RNA degradation. The degradation of *sodB* mRNA has been extensively studied in *E. coli* in absence and presence of RyhB (37, 38, 41, 57, 58). The decay depends on the RNA degradosome machinery, which comprises the single-strand specific RNase E, the 3’-5’ exoribonuclease PNPase, the RNA helicase RhlB and the enolase (reviewed in (59)). Degradation of mRNA in *E. coli* is often initiated by RNase E and subsequently followed by the attack of several exoribonucleases and oligoribonucleases to complete the degradation (60). Lifetimes of *E. coli* mRNAs can differ greatly since reported half-lives range from less than 1 min to 15 min or more (61, 62). Furthermore, mRNA lifetimes can be regulated in a translation-independent manner by binding of *trans*-acting regulatory factors such as sRNAs or RNA-binding proteins that impede or enhance RNase E cleavage (63–66). Together, these observations suggest that mRNA turnover is determined not by the number of cleavage sites but, rather, by the ease with which RNase E can gain access to them and the kinetics with which it cleaves. Interestingly, the association between the RNA degradosome and S1 (together with Rho) was observed in *Caulobacter crescentus* (which has S1 with 6 OB-fold domains) at low temperature (67). Moreover, in *Salmonella typhimurium*, a loss-of-function mutation in r-protein S1 lacking domain 6, was identified as a suppressor for a RNase E temperature-sensitive (TS) mutation that affects its mRNA turnover ability (68). The S1Δ6 and the RNase E TS mutant strains have complementary phenotypes since RNase E TS mutant is heat sensitive, while S1Δ6 is cold sensitive. Furthermore, RNase E TS mutant decreases general mRNA half-lives and this phenotype is restored in the double mutant. In other words, it is possible that the C-terminus of S1 could directly influence the kinetics of RNA target cleavage by RNase E (**Figure 5B**). We propose that S1 could have two roles: (1) to prepare the mRNA site for optimal cleavage through its RNA unwinding capacity, acting either on or outside the ribosome; and/or (2) to facilitate the recruitment or recycling of the RNA degradosome, since S1 is able to bind both to RNase E and PNPase (69). Even though we favor a direct role of S1 in mRNA decay, we cannot rule out that its action on the rapid depletion of mRNAs repressed by RyhB would result from an indirect effect. Indeed, *in vitro* degradation of *sodB* mRNA was slower with the immature ribosomes isolated from *rpsA*Δ56 strain than the fully matured ribosomes isolated from WT strain containing full length S1 or from which S1 was removed, while the isolated proteins show no effect. Hence, we speculate that the RNase E would be stably associated with the unprocessed ribosomes prepared from *rpsA*Δ56 mutant strain. The lack of free RNase E might in turn induce slower degradation of *sodB* mRNA in presence of RyhB (**Figure 5A, lower panel**).

Another unexpected result was the fact that the RyhB-mediated activation of *shiA* mRNA was strongly affected in the mutant *rpsA*Δ56 strain. It was previously shown that RyhB acts together with Hfq to favor the recruitment of the ribosome and the formation of the initiation complex, which in turn stabilizes the mRNA. In addition, the secondary structure of *shiA* mRNA revealed an unpaired AU-rich sequence upstream the SD sequence (45), which is an appropriate binding site for S1 (34) and for a ribosome standby site (70). In such a model, the ribosome in standby would easily relocate to form a productive complex as soon as the RBS become accessible (**Figure 5C**). Because S1 has unwinding properties, it would help to prepare the binding site for RyhB. Together with Hfq, RyhB would strongly stabilize the open form of the RBS for efficient translation and as the consequence the mRNA would be stabilized.

Another major phenotype that we have identified is a strong defect of motility of the mutant strain, as reflected by the repression of the synthesis of FliC (**Figure 1A**). This result could also be explained by the involvement of S1 in mRNA decay. Indeed, *fliC* belongs to the motility cascade activated by FlhDC (71), which is itself protected from degradation by CsrA (72). The activity of CsrA is modulated by CsrB sRNA, that is able to sequestrate this RNA-binding protein (73). According to the transcriptomic data, CsrB expression is higher in the mutant *rpsA*Δ56 strain. As the consequence, the levels of *flhDC* mRNA drops and the expression of FliC is subsequently decreased (for a review, (74). In addition, CsrD is responsible for modulating CsrB level in the cell by promoting its degradation (75), and this decay requires an additional factor.

The affinity of r-protein S1 for unpaired AU-rich sequences, its ability to melt weak secondary structure elements, and the existence of 6 OB-fold domains endow the protein with the ability to adapt its mechanism of action according to the RNA and/or protein substrates, generating a panel of cellular functions. Our work highlights a new function of S1 in sRNA-dependent regulation and in rRNA maturation showing that in *Enterobacteriaceae*, S1 is at the crossroad of many functions, all linked to RNA metabolism. Whether these functions are conserved in bacteria carrying a shorter version of S1 remained to be studied.

## Acknowledgements

We are thankful to Philippe Hammann and Johanna Chicher for the help in the proteomic analysis, to Béatrice Chane-Woon-Ming for the R script used to generate the volcano plots and to Isabelle Caldelari and David Lalaouna for fruitful discussions. This work was supported by the Centre National de la Recherche Scientifique (CNRS; PR), by the French National Research Agency ANR (ANR-16-CE11-0007-01 to [P.R.]). This work of the Interdisciplinary Thematic Institute IMCBio, as part of the ITI 2021-2028 program of the University of Strasbourg, CNRS and Inserm, was supported by IdEx Unistra (ANR-10-IDEX-0002) and by SFRI-STRAT’US project (20-SFRI-0012), and EUR IMCBio (IMCBio ANR-17-EURE-0023) under the framework of the French Investments for the Future Program. Work in the Massé lab has been supported by an operating grant from the Canadian Institutes of Health Research (CIHR) to EM. BFL and KJB were supported by a Wellcome Trust Investigator award (200873/Z/16/Z). Mass spectrometry instrumentation was funded by the University of Strasbourg, IdEx Equipement mi-lourd 2015 and labEx NetRNA [ANR-10-LABX-0036].

## Material & Methods

### Strains, plasmids and oligonucleotides

All strains and plasmids, which were constructed and used in this study, are described in Supplementary Information. The oligonucleotides sequences are given in Supplementary Information. The *lacZ* fusion described in **Figure 2D** has been performed as previously described (38) in the WT and *rpsA*Δ56 context.

### Proteomics analysis

Protein extraction has been performed on bacteria *rpsA*1, Δ6 or Δ56 grown in LB at 37°C under constant agitation until OD_600_ = 0,4. Label free spectral count analysis was performed in triplicate using nanoLC–MS/MS. Protein samples were precipitated with 0.1 M ammonium acetate in 100% methanol and the protein pellets were further digested with sequencing-grade trypsin (Promega). For the analysis involving the *rpsA*Δ6 mutant missing the last C-terminal OB fold domain of protein S1, the samples were analyzed on a TripleTOF5600 mass spectrometer coupled to an NanoLC-2DPlus ChiP system (Sciex). For the analysis involving the *rpsA*Δ56 mutant missing the last two C-terminal OB fold domains of protein S1, the samples were analyzed on a QExactivePlus mass spectrometer coupled to an EASY-nanoLC-1000 (Thermo-Fisher Scientific). Data were searched against the *E.coli* updated UniProtKB database (release 2020_05) with a decoy strategy. Peptides were identified with Mascot algorithm (version 2.6, Matrix Science, London, UK) and then imported into Proline 2.0 software (http://proline.profiproteomics.fr/). Proteins were validated with Mascot pretty rank equal to 1, and 1% FDR on both peptide spectrum matches (PSM score) and protein sets (Protein Set score). The total number of MS/MS fragmentation spectra was used to relatively quantify each protein between the WT and mutant conditions performed in three independent biological replicates. The statistical analysis based on spectral counts was performed using a homemade R package (IPinquiry4 under https://github.com/) using the quasi-likehood negative binomial model from edgeR (R v3.5.0). For each identified protein, an adjusted p-value corrected by Benjamini-Hochberg was calculated, as well as a protein fold-change (FC) (**Table S1**). The results are presented in a Volcano plot using protein log2 fold-changes and their corresponding adjusted log10P-values to highlight enriched proteins in both conditions (**Figure 1A** and **Figure S1C**). The mass spectrometric data were deposited to the ProteomeXchange Consortium via the PRIDE partner repository with the dataset identifier PXD023838.

### Preparation of RNAs

The bacteria were grown in LB at 37°C under constant agitation until DO600 = 0,4. When necessary, the 2,2’-dipyridyl was added at 250 μM (point referred at t=0 min) and when necessary FeSO_4_ 100 μM at t=5 min. Rifampicin was added at a final concentration of (300 μg/ml). Bacteria are harvested at different time points, pelleted and frozen at −80°C. The genome of the bacteria is checked by PCR using AK68 and KAV04 primers. RNAs were extracted according to the FastRNA Pro protocol (Qbiogene). RNA preparation for transcriptomics analysis was performed in biological duplicates.

### Transcriptomics analysis

RNA samples (1 μg) from biological duplicates of WT and *rpsA*Δ56 cultures were ribo-depleted (Ribo-Zero rRNA Removal Kit (Bacteria) Illumina) and cDNA libraries were prepared using the adapter ligation strategy by Vertis NGS service (Germany). RNA samples were fragmented with ultrasound (4 pulses of 30 sec at 4°C) followed by a treatment with antarctic phosphatase and re-phosphorylated with polynucleotide kinase (PNK). Afterwards, oligonucleotide adapters were ligated to the 5’ and 3’ ends. First-strand cDNA synthesis was performed using M-MLV reverse transcriptase and the 3’ adapter as primer. The resulting cDNAs were amplified with PCR using a high fidelity DNA polymerase. The primers used for PCR amplification were designed for TruSeq sequencing according to the instructions of Illumina. The cDNA was purified using the Agencourt AMPure XP kit (Beckman Coulter Genomics). The cDNAs have a size range of 200-500 bp. The libraries were either paired-end sequenced using 2×75 bp read length (Replica R2 samples) or single-end sequenced using 50 bp read length (Replica 1 samples) on an NextSeq 500 system (Illumina). RNA-seq analysis was performed according to (76). Reads were processed and aligned on *E. coli* genome (NCBI RefSeq Accession NC_000913.3) using the Galaxy platform (77). We used DEseq2 to estimate enrichment values (P-value < 0.05; Fold change (FC) > 2) (**Table S2**). Transcriptomics data are available in the GEO database with the accession code GSE166046.

### Northern Blot

After separation on agarose gels (1-2 %) containing 20 mM guanidine thiocyanate or on 8 % polyacrylamide-7 M urea gels, 20 μg or 5-10 μg of total RNA, respectively, was transferred onto Hybond-N+ or Hybond-XL membranes (Amersham Bioscience). Cross-linking was performed by UV (1200 J). For detection of transcripts, DIG-labeled RNA probes (prepared according to the protocol provided by Roche, Cat. No. 11 277 073 910) or radiolabeled DNA probes and RNA probes were used (**Table S5**). Each experiment was reproduced at least three times. For the determination of the half-lives of *sodB* mRNA in the two strains, quantification of the remaining mRNA at the different time points was done by ImageQuant TL software (GE Healthcare Life Sciences).

### Proteins extraction and Western Blot analysis

Protein extraction was performed using the following protocol. Cold TCA solution was added to cells (5% final concentration) and the mixture was placed on ice for 10 min. After precipitation (15,000 g, 10 min), the protein precipitate was washed with 80% acetone (twice). Western blot analysis was performed as previously reported (78). Proteins were resuspended in protein-loading gel electrophoresis buffer, followed by separation on SDS-PAGE gel and transfer to nitrocellulose membrane. Mouse monoclonal ANTI-FLAG® M2 antibody (Sigma) was used at a dilution of 1:1,000. IRDye 800CW-conjugated goat anti-rabbit secondary antibody (Li-Cor Biosciences, Lincoln, NE, USA) was used at a dilution of 1:15,000. Western blots were revealed on an Odyssey infrared imaging system (Li-Cor Biosiences), and quantification was performed using the Odyssey 3.0 software.

### β-galactosidase assays

Kinetics assays for β-galactosidase activity were performed as described previously using a SpectraMax 250 microtitre-plate reader (Molecular Devices, Sunnyvale, CA, USA) (45). Briefly, overnight bacterial culture incubated at 37°C were diluted 1,000-fold in 50 ml of fresh LB medium and grown with agitation (220 rpm) at 37°C. When required, expression of respective sRNAs was induced by addition of 0.1% arabinose at OD_600nm_ = 0.1. Specific β-galactosidase activities were calculated using the formula Vmax/OD_600nm_ when cells reached an OD_600nm =_ 0.5 -0.8 (exponential phase of growth). Data represent the mean of three independent experiments (± standard deviation, SD).

### Ribosome purification and Toeprinting

The preparation of the *E. coli* 70S, 30S subunits, and toeprints were performed as previously described (79) (see Supplementary Information). Toeprint was done on *sodB 119* mRNA using fluorescently labeled s*odB*rev2 primer. After primer extension with reverse transcriptase, the cDNA products were analyzed by capillary electrophoresis (3130x Genetic analyzer Applied Biosystems) and data processed using QuShape software (80). For experimental details, see the supplementary materials.

### *In vitro* translation assays

The PURExpress ΔRibosome (NEB #E3313) kit was used according to the commercial protocol in the presence of Met-S^35^ (https://www.neb.com/~/media/Catalog/All-Products/0D1F4E4BB3F14EFC9DF22C6463654CE4/Datacards%20or%20Manuals/manualE6800.pdf). The reaction mix has been reduced to 10 μL, *sodB*-FL mRNA was used at 0,4 μM (1 μg) in the presence of ribosomes 70S purified from *rpsA*1 or *rpsA*Δ56 strains at 2,4 μM, with increasing concentration of RyhB (2 and 4 μM). The reaction is incubated 2h at 37°C, loaded on a 15% SDS-PAGE colored with Coomassie blue and revealed by autoradiography. A specific band at the bottom of the gel was used to normalize the signal.

### Analysis of RNA-RNA binding *in vivo*

Affinity purification assays were performed as described in (78). The *E. coli* bacterial strains (WT and *rpsA*Δ56 strains) were grown to an OD_600nm_ of 0.4, at which point 0.1% arabinose was added to induce the expression of MS2-RyhB or RyhB during 10 min. Cells equivalent to 40 OD_600nm_ were chilled for 10 min on ice. RNAs were extracted following the hot-phenol protocol from 600 μL of culture (input). The remaining cells were then centrifuged, resuspended in 1mL of buffer A (20 mM Tris-HCl at pH 8.0, 150 mM KCl, 5 mM MgCl_2_, 1 mM DTT), and centrifuged again. Cells were resuspended in 2 mL of buffer A and lysed using a French Press (430 psi, three times). Lysate was then cleared by centrifugation (17,000 g, 30 min, 4°C). The soluble fraction was subjected to affinity chromatography at 4°C composed of 75 μL of amylose resin bound to 200 pmol of MS2-MBP protein in a Bio- Spin disposable chromatography columns (Bio-Rad). After washing, the cleared lysate was loaded onto the column, and washed with 5 mL of buffer A. RNA and proteins were eluted from the column with 1 mL of buffer A containing 15 mM maltose. Eluted RNA was extracted with phenol-chloroform, followed by ethanol (3 vol) precipitation of the aqueous phase in the presence of 20 mg of glycogen. For protein isolation, the organic phase was subjected to acetone precipitation. RNA samples were then analyzed by Northern blot and protein samples by Western blot.

### Motility track/soft agar

An overnight culture of the different strains was grown in LB at 37°C upon constant agitation and used the next day to inoculate a fresh day culture in LB at 37°C upon constant agitation to reach an OD_600nm_ = 0.4. Then, 2uL of culture was dropped onto a soft agar petri dish (tryptone 13 g/L, NaCl 7 g/L, agar-agar 0.3%) and grown overnight at 37°C. For motility tracking, the day culture was inspected between slide and slip cover using an optical microscope, and movies were acquired. These videos were processed using imageJ using mtrack2 plugin (1 out of 50 frames).

### Degradosome purification

The recombinant RNA degradosome was purified from *E. coli* as described in (81).

### *In vitro* kinetics of *sodB* degradation in presence of ribosomes and RyhB

The degradation *sodB* was monitored *in vitro* using the purified RNA degradosome under various conditions and as a function of time. The reactions were performed at 37°C from 5 to 20 min in a final volume of 6 μl containing the Degradosome Buffer (Tris HCl pH 7.5 25 mM, NH_4_Cl 50 mM, DTT 1 mM, KCl 50 mM, MgCl_2_ 10 mM, RNasine (Promega) 1 U/μl), the uniformly radiolabeled *sodB* mRNA (300 nM) free or bound to RyhB sRNA (2 μM), *E. coli* ribosomes (500 nM), *E. coli* initiator tRNA (Sigma; 2 μM) and the purified RNA degradosome (40 nM). In other experiments, the ribosomes were substituted by the purified proteins WT S1 or Δ56 (500 nM) (see Supplementary Information ^6^). The reactions were stopped by adding to 5 μl of reaction 5 μl of Stop Solution (Tris HCl pH 7.5 100 mM, EDTA 12.5 mM, NaCl 150 mM, SDS 1%, Proteinase K (Sigma) 2 mg/ml) and incubated 30 min at 46°C. Then, 6 μl of Urea Loading Buffer (urea 7 M, xylene cyanol 0.025 %, bromophenol blue 0.025 %) was added and the RNA fragments were fractionated on a polyacrylamide 8% (1/20)-urea 7M gel electrophoresis. Quantification of the full length mRNA was done by ImageQuant TL software (GE Healthcare Life Sciences).

